# Glycoproteome remodelling and granule-specific *N*-glycosylation accompany neutrophil granulopoiesis

**DOI:** 10.1101/2023.01.18.524318

**Authors:** Rebeca Kawahara, Julian Ugonotti, Sayantani Chatterjee, Harry C. Tjondro, Ian Loke, Benjamin L. Parker, Vignesh Venkatakrishnan, Regis Dieckmann, Zeynep Sumer-Bayraktar, Anna Karlsson-Bengtsson, Johan Bylund, Morten Thaysen-Andersen

## Abstract

Neutrophils store microbicidal glycoproteins in cytosolic granules to fight intruding pathogens, but their granule distribution and formation mechanism(s) during granulopoiesis remain unmapped. Herein, we perform comprehensive spatiotemporal *N*-glycoproteome profiling of isolated granule populations from blood-derived neutrophils and during their maturation from bone marrow-derived progenitors using glycomics-assisted glycoproteomics. Interestingly, the granules of resting neutrophils exhibited distinctive glycophenotypes including, most strikingly, peculiar highly truncated *N*-glycosylation in the azurophilic granules. Excitingly, proteomics and transcriptomics data from discrete myeloid progenitor stages revealed that profound glycoproteome remodelling underpins the promyelocytic-to-metamyelocyte transition and that remodelling is driven primarily by changes in protein expression and less by the glycosylation machinery. Notable exceptions were the oligosaccharyltransferase subunits responsible for initiation of *N*-glycoprotein biosynthesis that were strongly expressed in early myeloid progenitors correlating with high glycosylation efficiencies of the azurophilic granule proteins. Our study provides spatiotemporal insights into the complex neutrophil *N*-glycoproteome featuring an intriguing granule-specific *N*-glycosylation formed by dynamic remodelling during myeloid progenitor-to-neutrophil maturation.

**Key points:**

- Systems glycobiology reveals that profound *N*-glycoproteome remodelling accompanies early neutrophil granulopoiesis
- Precision glycoproteomics produces detailed cartography of neutrophils that exhibit site-, protein- and granule-specific N-glycosylation

## Introduction

Neutrophils are critical first-line responders to infection and form a potent line of innate immune defence against invading pathogens^1,2^. Circulating in their resting state in blood, neutrophils are equipped with an arsenal of prepacked cytosolic granules^3^. The carefully timed mobilisation of these intracellular granules released upon specific environmental cues enables neutrophils to extravasate and traffic to inflammatory sites to elicit an effective, timely and well-balanced response against invading pathogens, which they fight by means of phagocytosis, degranulation and neutrophil extracellular traps (NETs)^3,4^.

Distinctly different granule populations are formed during the week-long maturation of neutrophils (granulopoiesis) in the bone marrow^3,5^. Azurophil (Az) granules are formed during the early myeloblast-to-promyelocyte (MB-to-PM) maturation stages whilst specific (Sp) granules are formed during the PM-to-myelocyte/metamyelocyte (MC/MM) stages. Finally, gelatinase (Ge) granules form during the late MM-to-band cell (BC) maturation stages. Secretory (Se) vesicles, although not considered granules *per se,* form another type of membrane-enclosed intracellular compartments within neutrophils that arise from endocytosis and plasma membrane invagination during the final BC-to-mature neutrophil maturation stage^3^. Now well documented, the granule populations harbour distinct proteome signatures; the temporal granule formation during granulopoiesis is, through a mechanism known as ‘targeting-by-timing’, a recognised contributor to the granule-specific protein repertoires of neutrophils^6,7^.

Protein *N*-glycosylation, the covalent attachment of *N*-glycans to motif-restricted asparagine residues (Asn-X-Ser/Thr, X ≠ Pro), adds important structural and functional heterogeneity to proteins. Both the neutrophil cell surface and granule-resident glycoproteins are richly decorated with *N*-glycans that facilitate and modulate key immune and trafficking functions of neutrophils^4^. For example, neutrophil proteins are known to carry.*N*-glycans containing Lewis X (Le^x^) and sialyl-Lewis X (sLe^x^) glycoepitopes^8^, which interact with endothelial P- and E-selectins to mediate neutrophil tethering, rolling and extravasation to peripheral tissues^9,10^.

Following our discovery of paucimannosidic *N*-glycans (Man_1-3_GlcNAc_2_Fuc_0-1_) in the neutrophil-rich sputum from cystic fibrosis-affected individuals^11,12^, we and others have demonstrated that neutrophils indeed produce paucimannosylation that abundantly decorates microbicidal glycoproteins residing in the Az granules e.g. cathepsin G (CTSG), neutrophil elastase (ELANE), and myeloperoxidase (MPO)^12–18^. The biological roles of paucimannosidic proteins in neutrophils, however, remain poorly understood despite emerging evidence pointing to their involvement in key innate immune processes^19,20^.

While protein glycosylation is widely recognised to shape neutrophil function^4^, the spatial *N*-glycoproteome distribution across the neutrophil compartments and the dynamic glycoproteome changes (remodelling) that accompany the dramatic metamorphosis associated with the differentiation of myeloid progenitor cells to mature polymorphonuclear (PMN) blood cells remain unknown. Herein, we comprehensively profile the dynamic and complex neutrophil *N*-glycoproteome with spatial (granule) and temporal (maturation stage) resolution focusing specifically on characterising the luminal (soluble) granule fractions hosting most microbicidal glycoproteins. Using glycomics-assisted glycoproteomics of isolated neutrophil granules complemented with multi-omics data interrogation of myeloid progenitors still developing in the bone marrow, we provide spatiotemporal insights into previously unknown neutrophil *N*-glycosylation processes. We find that the neutrophil *N*-glycoproteome exhibits fascinating site-, protein- and granule-specific *N*-glycosylation features and undergoes profound remodelling during the early stages of neutrophil maturation in the bone marrow.

## Methods

### Donors, neutrophil isolation and granule separation

Neutrophils were isolated from buffy coats from healthy donors at The Blood Center, Sahlgrenska University Hospital, Gothenburg, Sweden. According to Swedish law on ethical conduct in human research, ethics approval was not needed because the buffy coats were provided anonymously. Resting neutrophils were isolated to >95% purity in four biological replicates each comprising pooled buffy coats from four individuals as described^14^. Neutrophil granules were separated using an established three-layered Percoll method^21^, monitored using established immunoblot and enzyme activity assays^14^ and validated using proteomics. Granules were lysed and the luminal (soluble) protein extract collected after ultracentrifugation. See **Extended methods** for details of all experimental procedures.

### Glycome profiling

*N*-glycans were prepared for glycomics as described^22^. Briefly, *N*-glycans were released using *N*-glycosidase F, hydroxylated and reduced prior to *N*-glycan desalting. The *N*-glycans were profiled in negative polarity mode on a Velos Pro linear ion trap connected to a Dionex Ultimate-3000 HPLC (Thermo Fisher Scientific) using an established PGC-LC-MS/MS method^14^. GlycoMod and GlycoWorkBench v2.1^23^ assisted the otherwise manual *de novo* glycan annotation as described^24^. Briefly, *N*-glycans were identified based on mass, MS/MS fragmentation pattern and relative/absolute PGC-LC retention time. AUC-based glycan quantification was performed using Skyline v.19.1^25,26^.

### Glycoproteome profiling

Proteins were precipitated, reduced, alkylated and digested using trypsin. Peptides were desalted using C18-SPE and either analysed directly by LC-MS/MS or enriched for glycopeptides using hydrophilic interaction liquid chromatography SPE as described^15^.

Unenriched peptides were analysed in positive ion polarity using a Q-Exactive HF-X Hybrid Quadrupole-Orbitrap coupled to an Easy nLC-1200 HPLC (Thermo Scientific). Peptides were separated using reversed-phase C18 chromatography and fragmented using higher-energy collisional dissociation (HCD)-MS/MS.

Enriched glycopeptides were analysed in positive ion polarity using an Orbitrap Fusion coupled to a Dionex 3500RS HPLC (Thermo Scientific). Glycopeptides were separated using reversed-phase C18 chromatography and fragmented using HCD-, ETD- and CID-MS/MS.

Employing the Andromeda search engine and the generated HCD-MS/MS data, MaxQuant v1.6.10.43^27^ was used for protein identification and quantification. Identifications were filtered to 1% protein FDR. Label-free AUC-based quantification was performed and protein abundance was calculated based on the normalised protein intensity (LFQ intensity)^28^.

Glycopeptides were identified and quantified from the HCD-MS/MS data using Byonic v4.5.2 (Protein Metrics Inc) in Proteome Discoverer v.2.5. Glycopeptide identifications were filtered to FDR <1%. CID- and ETD-MS/MS data were used to manually confirm glycopeptides. (Glyco)peptides were quantified based on AUC of precursor ions using the Minora Feature Detector or glycoPSMs.

### Interrogation of available proteomics and transcriptomics resources

Proteomics data of immature and mature neutrophils (PXD013785)^29^ were interrogated for intact glycopeptides (see above). Matching transcriptomics data (https://blueprint.haem.cam.ac.uk/neutrodiff)^30^ were interrogated for glycoenzymes (e.g. OST) and granule protein expression patterns.

### Statistical analyses

Significance was assessed using unpaired two-tailed Student’s t-tests or ANOVA followed by Tukey test for multiple comparison with FDR < 0.05. For transcriptomics and glycomics data analyses, significance was corrected for multiple test comparisons. *p* < 0.05 was considered significant. GraphPad Prism v9.4.1 (Dotmatics) and Perseus v2.0.7.0^31^ were used for statistical analyses. Correlation analyses were performed using Pearson correlation coefficient using Excel. PCA was performed using Metaboanalyst v. 5.0^32^.

### Data sharing statement

All LC-MS/MS data and metadata have been deposited in public repositories. Specifically, glyco/proteomics LC-MS/MS data were deposited to PRIDE (PXD039387, PXD021131). Glycomics LC-MS/MS raw data were deposited to GlycoPOST (GPST000315).

## Results

### Spatiotemporal glycoproteome profiling of resting and maturing neutrophils

Spatial *N*-glycoproteome profiling of resting neutrophils was performed by applying integrated glycomics-assisted glycoproteomics^33,34^ to isolated granule populations (**Figure 1A**). The temporal *N*-glycoproteome remodelling accompanying granulopoiesis was explored by interrogating robust proteomics and transcriptomics resources generated from discrete myeloid progenitor subsets^29,30^ for glycopeptide and glyco-enzyme expression patterns (**Figure 1B**).

**Figure 1.**
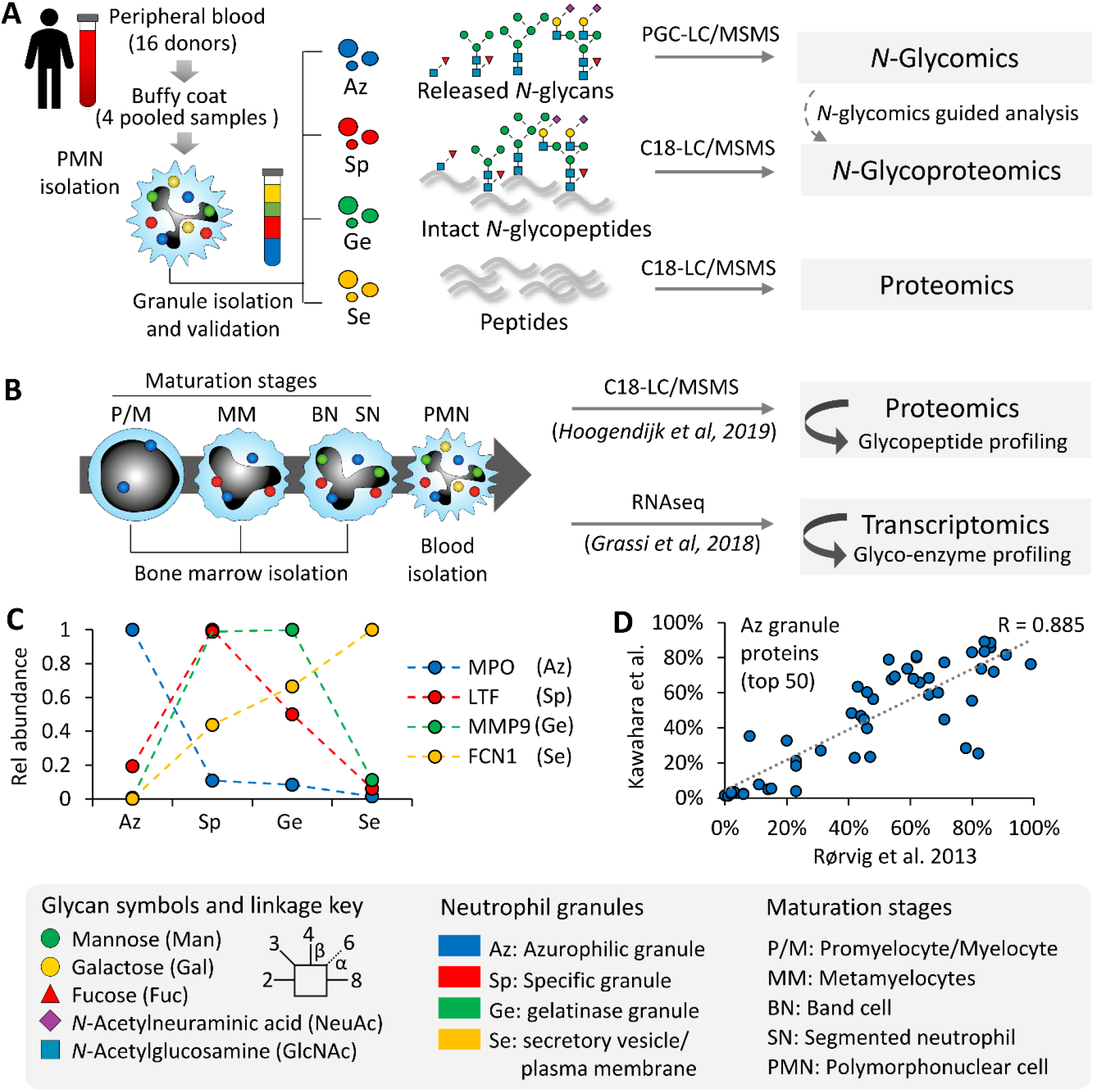
Spatiotemporal *N*-glycoproteome profiling of neutrophil granules and myeloid maturation stages using systems glycobiology approaches. **A)** Spatial *N*-glycoproteome profiling using quantitative glycomics-assisted glycoproteomics of the four major granule populations isolated from resting neutrophils. **B)** Temporal profiling of the *N*-glycoproteome and the underpinning glycosylation machinery of discrete neutrophil maturation stages from bone marrow-derived myeloid progenitor cells through interrogation of available proteomics and transcriptomics resources^29,30^. Neutrophil granule isolation was orthogonally validated using the generated proteomics data by **C)** plotting the expression profile of known neutrophil granule markers i.e. myeloperoxidase (MPO) for azurophilic granules (Az), lactoferrin (LTF) for specific granules (Sp), matrix metalloproteinase-9 (MMP9) for gelatinase granules (Ge) and ficolin-1 (FCN1) for the secretory vesicles/plasma membrane (Se) and **D)** concordance of the 50 most abundant proteins identified in the Az granules against a key reference study also performing quantitative proteomics of separated neutrophil granules^7^. See **Supplementary Table S1** for the correlation of other granule datasets. See insert for key to the glycan symbols, linkage nomenclature and the neutrophil granules and maturation stages.

Separation of four granule populations of neutrophils was guided by established immunoblot and enzyme activity assays against known granule markers across the Percoll density fractions^14,21^. The enrichment of distinct granule subsets was orthogonally validated using the acquired proteomics data by plotting the expression profile of protein markers known to be elevated in each granule population (**Figure 1C**). Further supporting the precision of the granule separation, concordance was observed between our granule proteomics data and a landmark study also performing quantitative proteomics on separated neutrophil granules^7^ as illustrated by high correlation (R = 0.886) of proteins identified in the Az granules (**Figure 1D**) and in other granules (Sp: 0.854; Ge: 0.610; Se: 0.890) (**Supplementary Table S1**).

### Distinct *N*-glycome signatures across the neutrophil granules

Firstly, deep quantitative glycomics performed using PGC-LC-MS/MS revealed a total of 71 protein-linked *N*-glycan structures spanning 43 *N*-glycan compositions across the neutrophil granules (**Supplementary Table S2** and **Supplementary Data S1**).

Principal component analysis (PCA) of the *N*-glycomics data indicated that the neutrophil granule populations carry distinct *N*-glycome signatures (**Figure 2A**). The Az granules appeared particularly well separated from the other granule populations based on the *N*-glycome data. With respect to the *N*-glycan type distribution, the Az granules displayed abundant paucimannosylation (~55% of the total Az *N*-glycome) (**Figure 2B**). The Sp granules displayed mainly sialofucosylated complex-type *N*-glycans (~75%) while oligomannosidic- and less decorated complex-type structures were characteristic *N*-glycan types of both the Ge and Se granules.

**Figure 2.**
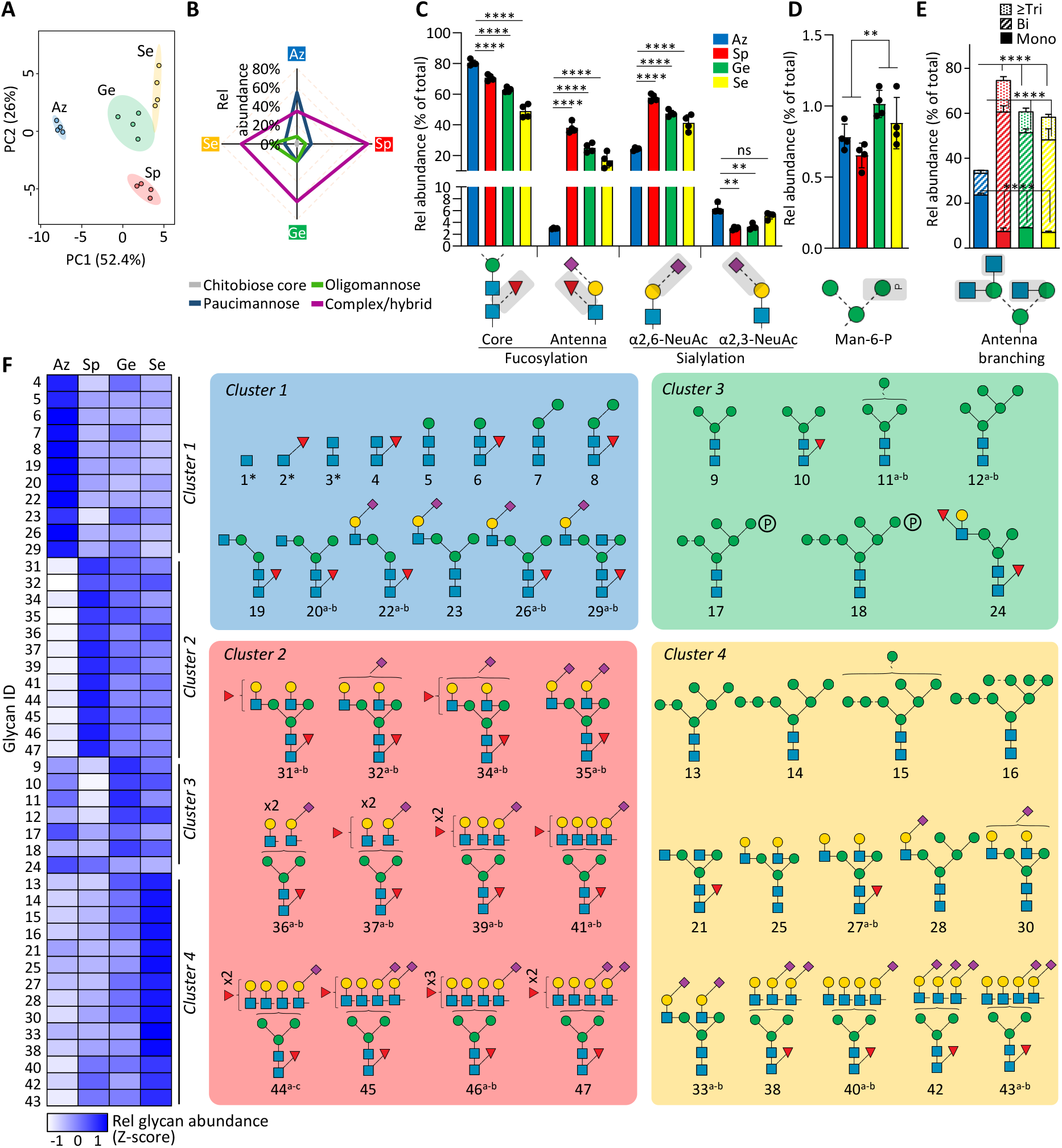
Granule-specific *N*-glycome signatures in resting neutrophils. **A)** PCA of quantitative *N*-glycomics data of the four granule populations (Az, Sp, Ge, Se) isolated from resting neutrophils. **B)** *N*-glycan class distribution across granules. Key glyco-features across granules including **C)** core/antenna fucosylation and α2,3-/α2,6-sialylation, **D)** mannose-6- phosphate (M6P) modifications and **E)** antennary branching patterns (relative abundance of each glyco-feature against the total *N*-glycome). For panel B-E, data plotted as mean and SD (n = 4 biological replicates, SD left out in panel B for simplicity, ANOVA test, \*\**p* < 0.01, \*\*\*\**p* < 0.0001, ns, not significant). Granule-specific differences were also observed for fucosylation, sialylation, M6P, and antennary branching when measured as a proportion of identified structures having the potential to be modified by the specific glyco-feature rather than against the total *N*-glycome, a more appropriate measure when exploring for biosynthetic relationships associated with observed *N*-glycome differences (**Supplementary Figure S1A- B**). **F)** Left: Unsupervised clustering (see **Supplementary Figure S1C** for dendrogram) and heatmap of *N*-glycomics data illustrating four major clusters representing the neutrophil granule populations. Right: *N*-glycan structures contributing to each cluster. The most common structure is depicted for glycan compositions exhibiting multiple isomers. *chitobiose core-type *N*-glycans reported at the glycopeptide level. See **Figure 1** for key. See **Supplementary Table S2** for glycan identifiers consistently used throughout the study.

Profound granule differences were also observed for the fine structural elements of the complex-type *N*-glycans. While core fucosylation (α1,6-fucosylation) was a prevalent feature across all granule subsets, the Sp, Ge and Se compartments exhibited significantly lower core fucosylation (~45-70%) compared to the Az granules (~80%, all *p* < 0.0001, ANOVA test) (**Figure 2C**). In contrast, antenna fucosylation forming key Lewis-type glycoepitopes was near-absent in the Az granules (~3%) while being prominent features in other granules (~20- 40%, all *p* < 0.0001, ANOVA test). Interestingly, sialyl linkage switching was also observed across the granule populations; the Az granules exhibited less α2,6-sialylation and high α2,3- sialylation relative to the other granules (all *p* < 0.01 except for Az vs Se α2,3-sialylation, ANOVA test). Albeit of generally low abundance (0.5-1% of *N*-glycome), mannose-6- phosphate (M6P) containing oligomannosidic *N*-glycans (M6-M7) were additionally found to be elevated in the Ge and Se granules relative to the Az and Sp granules (*p* < 0.01, student’s t-test) (**Figure 2D**). Lastly, dramatic differences in the antennary branching pattern were observed across the granules; unusual monoantennary complex-type *N*-glycans were features of the Az granules contrasting the bi-, tri- and higher branching patterns observed in the other granules (all *p* < 0.0001, ANOVA test) (**Figure 2E**).

Recapitulating the granule-specific *N*-glycome differences, unsupervised clustering analysis of the glycomics data indicated four major glycan clusters (cluster 1-4), one for each granule type, and revealed individual *N*-glycan structures contributing to the granule differences (**Figure 2D** and **Supplementary Figure S1C**). In short, cluster 1 (Az granules) was rich in peculiar chitobiose core-, paucimannosidic- and monoantennary complex-type *N*-glycans, cluster 2 (Sp granules) featured highly sialylated and core/antenna fucosylated complex-type branched *N*- glycans, cluster 3 (Ge granules) exhibited M6P-oligomannosylation, and cluster 4 (Se granules) oligomannose and less sialofucosylated complex-type *N*-glycans. Taken together, our comprehensive *N*-glycome map of luminal (soluble) proteins from the neutrophil granules, the most detailed to date, demonstrates prominent granule-specific *N*-glycan differences across the neutrophil granule subsets.

### Glycoproteomics reveals granule-, protein- and site-specific *N*-glycosylation in neutrophils

The glycoproteomics data of the neutrophil granules were searched for intact *N*-glycopeptides using a tailored glycan database comprising only *N*-glycans already identified by glycomics. Glycomics-guided glycoproteomics^22,33,34^ is beneficial as it reduces glycopeptide mis-identification and search time, two bottlenecks in glycoproteomics data analysis^35,36^, while it, in parallel, provides quantitative details of the *N*-glycans in the *N*-glycoproteome.

Collectively, a total of 27,663 *N*-glycoPSMs from 4,772 unique *N*-glycopeptides (unique protein, site and glycan) spanning 627 *N*-glycosites and mapping to 352 source *N*-glycoproteins were identified across the neutrophil granules (**Supplementary Figure S2A** and **Supplementary Table S3**). Despite using highly conservative confidence thresholds to ensure accurate glycopeptide identification, our analysis provides the, to date, deepest coverage of the neutrophil *N*-glycoproteome, greatly expanding on findings from a recent neutrophil glycoproteomics study performed without granule fractionation^13^ (**Supplementary Figure S2B-C**). Our glycomics-informed glycoproteomics approach therefore facilitated both an extensive glycoproteome coverage, and, importantly, an accurate and unbiased view of the neutrophil *N*-glycoproteome as supported by consistent correlations between our glycomics and glycoproteomics data across the investigated granule subsets (Az: R = 0.588, Sp: 0.505, Ge: 0.529, Se: 0.419, **Supplementary Figure S3A-B**). Importantly, the glycopeptides identified in the enriched and unenriched fractions showed a similar glycan distribution (Az: R = 0.875, Sp: 0.649, Ge: 0.789, Se: 0.906), supporting that both sample types enable accurate *N*-glycoproteome profiling of the neutrophil granules (**Supplementary Figure S3C-D**).

Interestingly, the glycopeptide spectral prevalence in the Az granules (averaging 2,558 *N*- glycoPSMs/run) far outnumbered the glycopeptide spectral count in other granules (700-1,501 *N*-glycoPSMs/run) (**Figure 3A**). Despite the higher proportion of glycopeptide spectra in the Az datasets, a similar number of unique glycopeptides but mapping to less source *N*- glycoproteins (94 proteins) was observed for the Az granules relative to the other granule populations (139-199 proteins/granule type), suggesting a higher glycan density per protein and, therefore, a higher occupancy of the available glycosylation sites (higher glycosylation ‘efficiency’) of the proteins residing in the Az granules.

**Figure 3.**
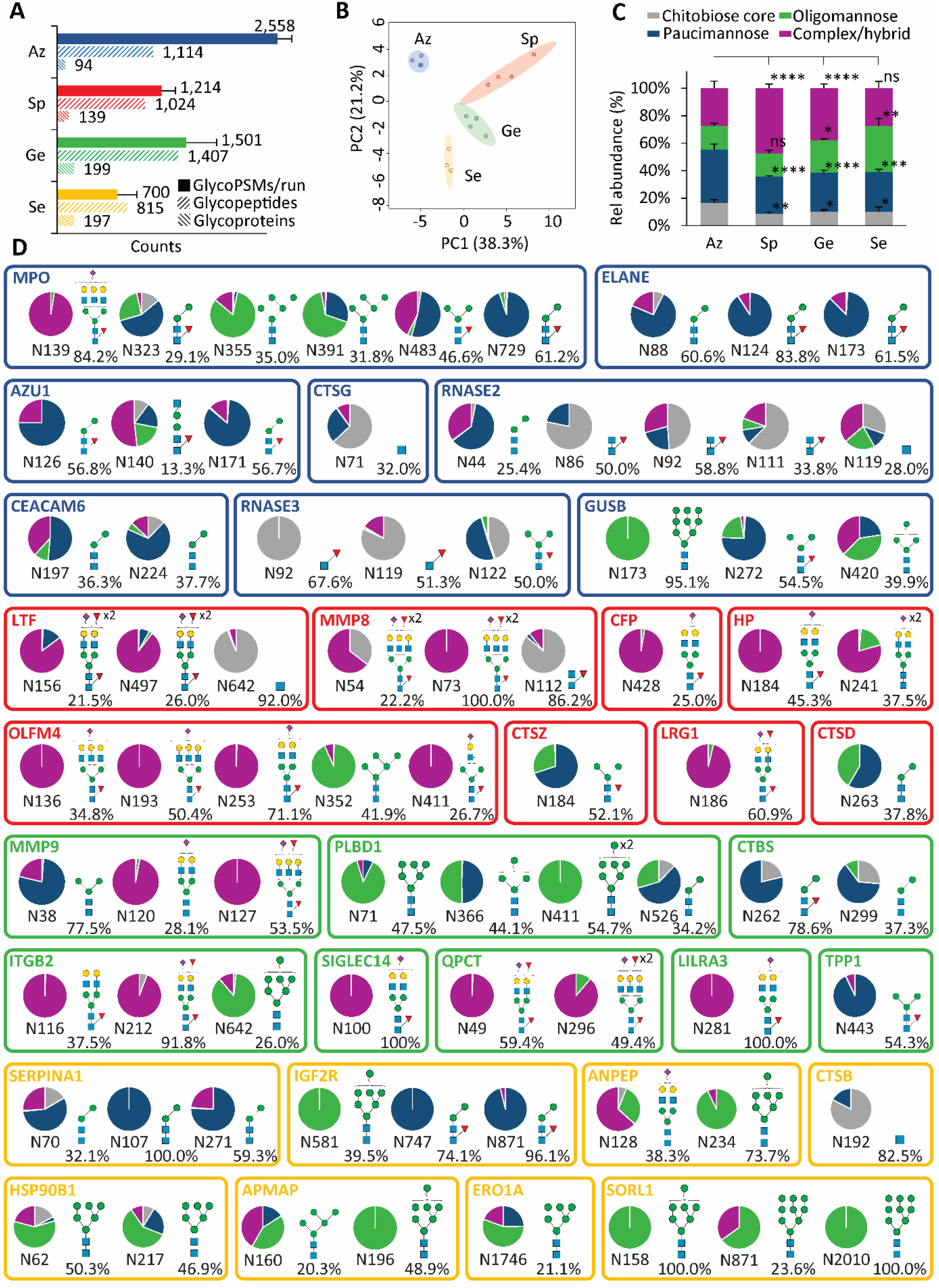
Glycoproteomics reveals site-, protein- and granule-specific *N*-glycosylation across the neutrophil granules. **A)** Glycoproteome coverage across the neutrophil granules as measured by *N*-glycoPSM counts, unique *N*-glycopeptides (unique protein, site, glycan), and source *N*-glycoproteins identified by quantitative glycoproteomics. GlycoPSM data are plotted as mean ± SD while glycopeptide/glycoprotein data are plotted as total counts from n = 4 biological replicates. **B)** Glycoproteomics-informed PCA of the neutrophil granules (Az, Sp, Ge, Se). **C)** *N*-glycan type distribution across neutrophil granules. Data plotted as mean ± SD (n = 4 biological replicates, \**p* < 0.05, \*\**p* < 0.01, \*\*\*\**p* < 0.0001, ns, not significant, ANOVA test). **D)** Extensive *N*-glycan diversity across sites and proteins as illustrated for the eight most abundant *N*-glycoproteins (out of 100-200 glycoproteins/granule) across each neutrophil granule. The most abundant glycoform (and its relative abundance) is indicated for each site. See **Figure 1** for key.

PCA of the glycoproteomics data recapitulated the glycomics-informed separation of the four granule populations supporting their distinctive glycophenotypes and highlighted again the unique glycosylation features of the Az granules (**Figure 3B**). In line with the glycomics data, the glycoproteomics data indeed showed that the Az granules exhibit higher levels of chitobiose core- and paucimannosidic-type *N*-glycans and lower levels of oligomannosidic- and complex-type *N*-glycans relative to the other compartments (**Figure 3C**).

The information-rich glycoproteomics data revealed a profound microheterogeneity featuring an extensive diversity and variation of the *N*-glycan distribution across sites and proteins as illustrated for the eight most abundant glycoproteins (of ~100-200 glycoproteins/granule) in each granule (**Figure 3D** and **Supplementary Table S4**). Importantly, while each glycoprotein (and each site) exhibited distinct glycosylation patterns, the granule-specific *N*-glycosylation features documented by glycomics were recapitulated in the glycoproteomics data as illustrated by similar glycan-based clustering patterns across granules (**Supplementary Figure S4**).

Taken together, our comprehensive glycomics-assisted glycoproteomics analysis of the four major granule populations of resting neutrophils provides deep spatial insight into the neutrophil *N*-glycoproteome that exhibits fascinating granule-, protein- and site-specific *N*- glycosylation features.

### Profound glycoproteome remodelling during neutrophil granulopoiesis

To begin to unpick mechanisms driving the intriguing granule-specific *N*-glycosylation in resting neutrophils, we investigated longitudinally the *N*-glycoproteome remodelling accompanying the granulopoiesis process in the bone marrow. Specifically, we mined robust proteomics and transcriptomics data collected from discrete myeloid progenitor stages for previously overlooked *N*-glycopeptide and glyco-enzyme expression patterns^29,30^. Owing to the high data quality, a total of 11,928 *N*-glycoPSMs from 2,276 unique *N*-glycopeptides mapping to 1,079 source *N*-glycoproteins (**Supplementary Figure S2A** and **Supplementary Table S5-S6**) and 45 glyco-enzymes driving diverse *N*-glycosylation biosynthetic processes (**Supplementary Table S7**) could be longitudinally profiled from these available resources.

In early granulopoiesis (P/M stage), the myeloid progenitors displayed prominent paucimannosylation (~45%) and chitobiose core-type (~25%) *N*-glycosylation, which were dramatically reduced in the subsequent maturation stage (MM stage, ~20% and ~15%, respectively), which instead featured elevated complex- (~15%->35%) and oligomannosidic-type (~15%->30%) *N*-glycans (**Figure 4A** and **Supplementary Table S5**). The glycoproteome remodelling was notably reduced in mid- (MM) and late- (BN, SN) stage granulopoiesis.

**Figure 4.**
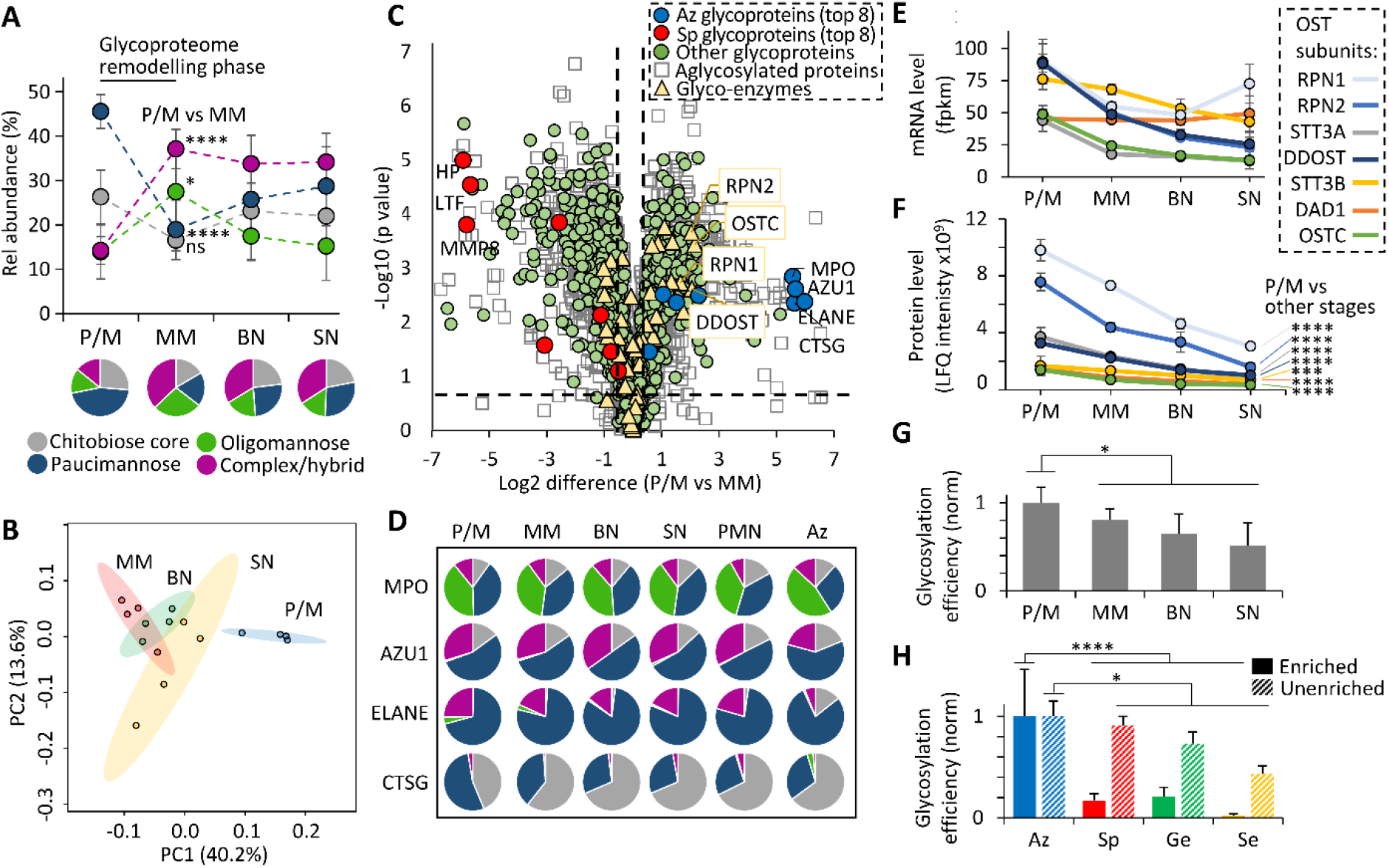
Dramatic glycoproteome remodelling in early granulopoiesis is accompanied by changes in proteome expression and the glycosylation initiation machinery. **A)** *N*-glycan class distribution during granulopoiesis as established by glycopeptide profiling from proteomics of key myeloid progenitor cell types (P/M, MM, BN, SN)^29^. **B)** *N*-glycopeptide- informed PCA of the four populations of myeloid progenitors. **C)** Differential transcript expression between P/M and MM progenitors. Permutation-based FDRs < 0.05 and S0 > 0.1 were calculated using Perseus. Dashed x/y lines represent significance thresholds. Proteins were assigned with granule location/glycosylation status based on glycoproteomics data (**Supplementary Tables S4**). Key *N*-glycosylation enzymes (yellow) including OST subunits (boxed, yellow) and the eight most abundant Az (blue) and Sp (red) glycoproteins amongst other glycosylated (green) and non-glycosylated (white) proteins are highlighted. **D)** Uniform *N*-glycosylation of key Az glycoproteins during granulopoiesis, in blood neutrophils (PMN) and in their principal granule location (‘Az’) of resting neutrophils. For simplicity, multiple sites were combined for the glycoprofile of MPO, AU1 and ELANE. **E)** Transcript (mRNA) and **F)** protein expression profile of the OST subunits during myeloid differentiation. Data plotted as mean ± SD (n = 4 biological replicates, \*\*\**p* < 0.001, \*\*\*\**p* < 0.0001, student’s t-tests, P/M vs other stages combined). Global protein *N*-glycosylation efficiency **G)** during granulopoiesis and **H)** across the corresponding neutrophil granules mapped separately for enriched (full bars) and unenriched (broken bars) glycoproteomics data. Data plotted as mean ± SD (n = 4 biological replicates. \**p* < 0.05, \*\*\*\**p* < 0.0001, student’s t-tests, P/M vs other stages combined. See **Figure 1** for key.

Early myeloid progenitors (P/M stage) indeed showed a distinctly different glycophenotype compared to progenitors in later maturation stages as illustrated by the separation of P/M cells by PCA, further supporting that strong *N*-glycoproteome remodelling accompanies early neutrophil maturation (**Figure 4B**). We therefore used transcriptomics to monitor the relative glycoprotein and glyco-enzyme expression during the P/M-to-MM transition (**Figure 4C**). Interestingly, the analysis revealed that prominent changes in the proteome expression as opposed to the glyco-enzymes accompany the *N*-glycoproteome remodelling phase. Expectedly, the Az proteins expressed at the P/M stage (e.g. MPO, AZU1, ELANE) and Sp proteins expressed at the MM stage (e.g. HP, LTF, MMP8) showed strong and opposite expression differences (up to 5-30 fold) between the two stages. In fact, ~75% of the transcriptome dataset (2,579/3,484 transcripts) demonstrated expression differences between the P/M and MM stages (Student’s t-test, FDR <0.05, **Supplementary Table S7**).

In line with the prominent protein expression differences and comparably minor changes to the glycosylation machinery during granulopoiesis, the glycosylation of key proteins enriched in the Az granules but found throughout all granule populations (MPO, AZU1, ELANE, CTSG) remained relatively unchanged during neutrophil maturation (**Figure 4D** and **Supplementary Table S5**). Taken together, these results indicate that the granule-specific *N*-glycosylation in resting neutrophils is driven primarily by glycoprotein repertoire changes during granulopoiesis as opposed to prominent global changes in the cellular glycosylation machinery. As a notable exception, the global expression analysis indicated altered P/M-to-MM expression of the oligosaccharide transferase (OST) enzyme complex responsible for the initiation of *N*- glycoprotein biosynthesis. We therefore used already mentioned -omics resources^29,30^ to monitor the transcript- and protein-level expression profile of the subunits forming the OST enzyme complex (STT3A/B, RPN1/2, DDOST, DAD1, OSTC) during granulopoiesis and found considerably reduced OST mRNA (**Figure 4E**) and protein expression (all *p* < 0.001, student’s t-test, P/M vs other stages combined) (**Figure 4F**) over the course of neutrophil maturation. Correlating with the observed OST dynamics, proteins expressed at the P/M stage and those identified in the corresponding Az granules displayed greater *N*-glycan occupancy (higher glycosylation efficiency) compared to proteins expressed later during differentiation and those residing in other corresponding granules (all *p* < 0.05, student’s t-test, P/M/Az vs other maturation stages/granules combined) (**Figure 4G-H**). Collectively, the temporal profiling of the *N*-glycoproteome and the *N*-glycosylation machinery during myeloid differentiation, the first of its kind, have revealed that profound glycoproteome remodelling accompanies early granulopoiesis and provides important system-wide molecular-level clues to mechanisms driving the granule-specific *N*-glycosylation observed in circulating blood neutrophils.

## Discussion

Neutrophils store cytosolic granules packed with highly glycosylated and potent hydrolytic enzymes, membrane receptors and microbicidal glycoproteins that can be mobilised in a timely and coordinated manner to allow neutrophils to traffic to sites of infection and fight invading pathogens^37^. While literature have documented diverse roles for protein *N*-glycosylation in neutrophil function^4^, the structural diversity and spatial distribution of the *N*-glycoproteome across the neutrophil granules and the temporal dynamics during the profound metamorphosis underpinning neutrophil granulopoiesis remain unmapped. Closing this knowledge gap is important since while under normal physiology and homeostasis neutrophils play protective roles throughout the body, it is becoming increasingly apparent that neutrophils, under certain conditions, play disease-promoting roles upon dysregulation of processes directly or indirectly involving protein glycosylation including impaired neutrophil maturation, recruitment, activation and turnover^4,38,39^.

In this study, we employed an integrated glycomics-assisted glycoproteomics method^22,34^ to comprehensively profile the *N*-glycoproteome of four major granule populations from resting neutrophils (Az, Sp, Ge, Se). The initial *N*-glycome profiling facilitated a quantitative view of the glycan fine structures and aided the informative (yet challenging^35,36^) *N*-glycoproteomics data analysis enabling us to establish with high precision the, to date, most detailed quantitative map of the heterogeneous *N*-glycoproteome across the neutrophil granules. Further, deep mining of *N*-glycopeptide and glyco-enzyme expression patterns using robust proteomics^29^ and transcriptomics^30^ resources obtained from discrete myeloid progenitor populations unveiled, with temporal resolution, the dynamic proteome expression and non-template-driven glycosylation processes accompanying neutrophil granulopoiesis.

Expanding significantly on previous efforts aiming to globally map neutrophil *N*-glycosylation ^12–14,40^, we here provide direct evidence for the distinctive glycophenotypes exhibited by the neutrophil granules and report on site-specific *N*-glycan features and carrier protein repertoires contributing to the intriguing granule-specific *N*-glycosylation observed in blood neutrophils. The Az granules, which consistently displayed the most distinctive *N*-glycosylation signatures, exhibited elevated levels of unconventional paucimannosidic-, chitobiose core- and monoantennary complex-type *N*-glycans. Recapitulating observations from targeted protein structure-function studies^12,15,17,18^, these unusual *N*-glycosylation features were found to be carried by MPO, ELANE and CTSG amongst ~100 other less studied Az glycoproteins. The discovery of peculiar *N*-glycosylation in the Az granules is important as this compartment is regarded as the microbicidal granule that neutrophils mobilise both intra- and extracellularly when combating pathogens at infection sites^3,37^. While the observation that unconventional *N*- glycans decorate a diverse repertoire of potent Az proteins is intriguing, their role(s) in shaping the microbicidal potential of the Az granules remains largely unstudied^4,20^. Our report of a bacteriostatic potential of the paucimannosidic *N*-glycans carried by ELANE towards clinical strains of *Pseudomonas aeruginosa* is, to our knowledge, the only literature on this topic^15^.

In addition to phagosomal fusion, a subset of the Az granules degranulate their content into the extracellular environment upon neutrophil activation^41^. Thus, there is also the potential for the unconventional glycosylation of the Az proteins to mediate cell-cell communication through putative signalling receptors towards paucimannose^19^ and induce production of autoantibodies in neutrophil-related auto-immune diseases, such as anti-neutrophil cytoplasmic autoantibody-associated vasculitis^18^. Supporting the former, we and others have demonstrated that mannan-binding lectin (MBL2)^12^ and the macrophage mannose receptor (CD206)^42^ display affinities to paucimannosidic glycoepitopes, but lectin receptors with high affinity/avidity to paucimannose and any cell-cell communication and other functional consequences resulting from their interaction require further exploration. Since neutrophil-derived MPO reportedly is internalised via endocytosis by CD206 on macrophage cell surfaces^43^, paucimannosylation may proposedly also function as danger associated molecular patterns^44^ or alternatively serve as ‘removal tags’ of the potent (thus potentially harmful) Az proteins to prevent damage to host tissues following excessive release and pathogen killing.

Our finding that biologically-relevant glycoepitopes (sLe/Le) decorate a considerable subset of the ~140 glycoproteins e.g. LTF, MMP8, and LRG1 identified in the Sp granules is in line with (but considerably expands on) existing literature^14,45,46^. Lewis-type epitopes are known to mediate neutrophil tethering and rolling, and, in turn, facilitate extravasation to inflammatory sites through interactions with endothelial selectins^47^. However, literature points primarily to membrane-embedded glycoproteins residing in readily mobile secretory vesicles e.g. PSGL-1, ESL-1, and CD44 being responsible for this trafficking process^4^. Our study reveals that many luminal Sp glycoproteins also carry s/Le glycoepitopes. The extracellular roles of these glycoproteins remain essentially unstudied apart from a few reports suggesting that Le^x^ on luminal Sp glycoproteins (e.g. LTF) mediates binding to the scavenger receptor C-type lectin (SRCL) expressed on endothelial cells to facilitate internalisation and clearance of such bioactive (and potentially harmful) glycoproteins^45,48^.

Finally, the Ge and Se granules each hosted ~200 *N*-glycoproteins carrying predominantly oligomannosidic- and less sialylated/fucosylated *N*-glycans. While traditionally regarded as an intracellular glyco-feature pertaining to incompletely processed *N*-glycoproteins trafficking the secretory machinery, oligomannosylation is increasingly reported i) on mature (fully processed) neutrophil glycoproteins^4,19^, ii) to decorate the surface of neutrophils^49^ and other blood^50^ and epithelial^51,52^ cells, and iii) to be elevated in cancer^53,54^ and other neutrophil-related diseases^55^. Our study supports that oligomannosylation is a prominent feature of the Ge and Se compartments. This is of potential significance since neutrophils readily expose, in a context-dependent manner, their granule content from these mobile compartments to the extracellular environment to enable communication with surrounding tissues or removal from circulation^4,19^.

Excitingly, our investigation of myeloid progenitors in different maturation stages revealed considerable plasticity in key *N*-glycosylation processes and detailed a global remodelling of the *N*-glycoproteome during neutrophil granulopoiesis. Particularly dramatic *N*-glycoproteome remodelling was found to accompany early maturation (P/M-to-MM transition). Notably, the Az and Sp granules formed during the P/M and MM differentiation stages, respectively, exhibited striking glycoproteome differences providing important mechanistic clues to the formation of the distinct glycoproteome characteristics across the granule populations. The P/M-to-MM transition is important as it marks the change from self-replicating promyelocytes to non-dividing and committed metamyelocytes, a transition that is accompanied with notable phenotypic (morphological, molecular, functional) alterations^29^. This is, to the best of our knowledge, the first report to demonstrate glycoproteome remodelling in developing myeloid progenitor cells. In fact, maturation stage-specific glycosylation appears scarcely investigated with recently available and highly informative glycoproteomics approaches. Examples from the glycobiological literature include our own report on dynamic glycosylation underpinning myogenesis and muscle development^56^, and *N*-glycoproteome changes during neuronal^57^, and cardiomyocyte^58^ differentiation reported by other groups.

Importantly, our multi-omics interrogations established that the *N*-glycoproteome remodelling occurring during granulopoiesis is driven primarily by temporal changes in the proteome expression patterns leading to different glycoprotein repertoires across the individual granule populations. Less prominent were changes within the glycosylation machinery responsible for the enzymatic processing of nascent *N*-glycoproteins trafficking the secretory pathway. This observation is important as it illustrates alternative factors (other than the glycosylation machinery) mediating glycoproteome remodelling in neutrophils, and serves to explain how some *N*-glycoproteins expressed throughout granulopoiesis (e.g. MPO, ELANE) can receive relatively uniform glycosylation regardless of the timing of their expression and their granule location (Figure 4D). To this end, both the protein repertoire and the glycosylation machinery can be considered determinants for the distinct glycophenotypes of the neutrophil granules. However, beyond relatively simple solvent accessibility><glycan processing relationships^59,60^, a molecular level understanding of how neutrophil proteins carry information that enable their individualised modification patterns across the neutrophil granules is still lacking. Finally, we acknowledge that other dynamic factors such as Golgi restructuring, altered micro-trafficking routes and changes to non-enzymatic components of the glycosylation machinery (i.e. nucleotide-sugar levels and transporters), all of which were not explored in this study, may also accompany granulopoiesis consequently adding another level of complexity to consider when attempting to decode mechanisms governing the dynamics of the neutrophil glycoproteome. As a notable exception to the relatively stable glycosylation machinery during neutrophil granulopoiesis, expression of the OST enzyme complex responsible for the initiation of *N*- glycoprotein biosynthesis was significantly reduced during neutrophil maturation, which correlated with a relatively high glycosylation efficiency of proteins in the Az granules relative to other granules (Figure 4E-H). We are not aware of similar reports of OST dynamics during the maturation of neutrophils (or other cell types), but a growing body of evidence points to context-dependent OST regulation and interesting links to cancer, immune evasion and ER stress^61,62^. It is tempting to speculate that Az proteins require high glycosylation occupancy to prevent premature proteolytic degradation during the extended storage in these highly hydrolytic compartments hosting many potent proteases (e.g. ELANE, CTSG). Although energetically expensive, the installation of voluminous and flexible glycans on protein surfaces is in fact shown confer effective protection against proteolysis^63,64^.

In steady-state conditions, neutrophils form from haematopoietic stem and progenitor cells in the bone marrow^1^. However, during systemic infection and inflammation involving a prolonged demand for neutrophils (e.g. cancer and chronic infections), the haematopoietic system switches from steady-state to ‘emergency granulopoiesis’^65,66^. This so-called ‘left-shift’ results in the appearance of neutrophil precursors in circulation featuring immature phenotypes including aberrant (progenitor-like) morphology and, presumably, an incomplete repertoire of granules and/or granule content. Considering our discovery of maturation stage-specific glycosylation, we hypothesise that immature neutrophils arising from emergency granulopoiesis display an “immature” (not fully remodelled) *N*-glycoproteome^4^. Impaired granulopoiesis is also central to acute myeloid leukemia (AML)^67^, in which morphologically immature blood neutrophils, expectedly exhibiting aberrant glycosylation, were found to show decreased capacity for NET formation^68^. The link (if any) between poor NET formation and aberrant glycosylation of AML neutrophils remains unexplored. Guided by our high-precision *N*-glycoproteome map of healthy neutrophils, future glycoprofiling efforts of immature neutrophils from individuals exhibiting emergency granulopoiesis and AML patients are required to substantiate these speculations and establish causal links to neutrophil dysfunction. In conclusion, our detailed spatiotemporal *N*-glycoproteome profiling of developing myeloid progenitors and mature neutrophils has painted the, to date, most comprehensive picture of the intriguingly complex neutrophil *N*-glycoproteome and has laid an important foundation to explain its formation. Our systems glycobiology (multi-omics) approach revealed that strong *N*-glycoproteome remodelling underpins neutrophil granulopoiesis, which, when considered with the widely accepted ‘targeting-by-timing’ model^6,7^, provides new mechanistic clues to the distinctive glycophenotypes exhibited by each neutrophil granule population as detailed herein. Importantly, our data-rich omics-centric study serves as a valuable resource to guide future explorations targeting the biological roles and dysregulation of neutrophil glycosylation that holds an untapped clinical potential for the discovery of glycoprotein-related markers for and therapeutics against a spectrum of neutrophil-related disorders.

## Acknowledgments

RK is supported by the Cancer Institute of New South Wales (ECF181259). JU is supported by a Macquarie Research Excellence Scholarship. SC was supported by an International Macquarie Research Excellence Scholarship (iMQRES 2017152). IL was supported by an International Research Training Program Scholarship funded by the Australian Government. HCT was supported by a Macquarie University COVID-19 fellowship. ZSB was supported by a JSPS fellowship. JB was supported by grants from the Swedish Research Council (2019-01123) and TUA Research Funding; The Sahlgrenska Academy at University of Gothenburg/Region Västra Götaland, Sweden (TUAGBG-917531). AKB is supported by the Swedish Research Council (2018-03077). MTA is supported by an Australian Research Council Future Fellowship (FT210100455).

## Author contributions

RK, JU, IL and MTA designed experiments. RK, SC, HT, IL, BLP and VV conducted experiments. RK, RD, ZSB, AKB, JB and MTA established laboratory methodologies. RK, JU, SC, HT, IL, ZSB and MTA analysed the data. BLP, VV, RD, AKB and JB provided reagents for this study. RK, JU and MTA wrote the manuscript. RK and MTA supervised this study and acquired funding. All authors have reviewed and approved the manuscript.

## Disclosure of conflicts of interests

The authors declare no competing interests.

## Notes

### Competing Interest Statement

The authors have declared no competing interest.

### Summary of Updates

Manuscript updated (shortened) in preparation for journal submission

