## Extended Methods and Supplementary Figures for "Glycoproteome remodelling and granule-specific *N*-glycosylation accompany neutrophil granulopoiesis"

**For**

[1]School of Natural Sciences, Macquarie University, Sydney, NSW, Australia; [2]Cordlife Group Limited, Singapore, Singapore; [3]Department of Anatomy and Physiology, University of Melbourne, Melbourne, VIC, Australia; [4]Department of Rheumatology and Inflammation Research, Institute of Medicine, Sahlgrenska Academy, University of Gothenburg, Gothenburg, Sweden; [5]Department of Life Sciences, Chalmers University of Technology, Gothenburg, Sweden; [6]Department of Oral Microbiology and Immunology, Institute of Odontology, Sahlgrenska Academy, University of Gothenburg, Gothenburg, Sweden; [7]Institute for Glyco-core Research (iGCORE), Nagoya University, Nagoya, Japan

R.K. and J.U. contributed equally to this study.

Running title: Spatiotemporal glycoproteome remodelling in neutrophils

*Corresponding author
Associate Professor Morten Thaysen-Andersen, PhD
School of Natural Sciences
Macquarie University
NSW-2109, Sydney, Australia

### Extended Methods

#### Donors and neutrophil isolation

Buffy coats (~60 ml) from sixteen healthy individuals without any apparent health issues were obtained from The Blood Center, Sahlgrenska University Hospital, Gothenburg, Sweden. According to Swedish law on ethical conduct in human research, ethics approval was not needed because the buffy coats were provided anonymously and could not be traced back to a specific individual. For this reason, information of age and gender is not available. From these samples, four biological replicates (named donor v-z for sample/data labelling purposes) each comprising of a pool of buffy coats from four individuals were investigated. Resting neutrophils were isolated from the pooled buffy coats by removing firstly erythrocytes by dextran (1%) sedimentation (1 x g) and secondly monocytes and lymphocytes by centrifugation on a Ficoll–Paque density gradient. Any remaining erythrocytes were eliminated by hypotonic lysis in a Krebs–Ringer phosphate buffer containing 10 mM glucose and 1.5 mM $Mg^{2+}$ yielding a population of resting neutrophils with >95% purity as assessed by flow cytometry.

#### Neutrophil granule isolation

Neutrophil granules were separated using an established three-layered Percoll method[1]. Briefly, ~$10^9$ neutrophils/sample were treated with 8 mM diisopropylfluorophosphate (final concentration), resuspended in 10 ml homogenisation buffer containing 100 mM KCl, 3 mM NaCl, 3.5 mM $MgCl_2$, 10 mM PIPES (pH 7.4) with 1 mM $ATP(Na)_2$, 0.5 mM phenylmethylsulfonyl fluoride and gently disrupted in a nitrogen bomb (Parr Instruments, Moline, IL, USA) at 400 psi. A density gradient generated by 1.12 g/mL, 1.09 g/mL, and 1.05 g/mL Percoll (GE17-0891-01, GE Healthcare, Sweden) was used to separate the azurophilic (Az) granules, specific (Sp) granules, and gelatinase (Ge) granules and a combined fraction containing the secretory vesicles and the plasma membrane (Se) after centrifugation at 37,000 x g, 4°C, 30 min. The separation of these four neutrophil granules (designed a-d for sample/data labelling purposes) was guided by monitoring known granule markers using established immunoblot and enzyme activity assays across the density fractions as previously demonstrated[2] and orthogonally validated using the acquired proteomics data to map proteins known to be enriched in each compartment.

The pelleted granules were resuspended in the homogenisation buffer (above) and lysed using a tip sonicator at 1/3 power (8 MHz) three times for 30 s on ice. The luminal (soluble) proteins were separated from the membrane (insoluble) proteins of each granule fraction by ultracentrifugation at 100,000 x g, 4°C, 90 min. The supernatants containing luminal proteins were carefully collected and the protein concentrations determined using Bradford assays. The protein samples were stored at a concentration of 4–15 µg/µl at –20°C until further handling.

**Glycome profiling**

*N-glycan release and clean-up*

The luminal protein extracts from the neutrophil granules were reduced using 10 mM aqueous dithiothreitol (DTT) in 100 mM $NH_4HCO_3$ (pH 8.4), 45 min, 56°C, and then carbamidomethylated using 25 mM aqueous iodoacetamide (IAA) in 100 mM $NH_4HCO_3$ (pH 8.4), 30 min, 20°C, in the dark. The alkylation reactions were quenched using 25 mM aqueous DTT (all final concentrations). The *N*-glycans were released from the protein extracts and prepared for glycomics as described[3]. For each granule sample, ~20 µg total protein was blotted on a primed 0.45 mm polyvinylidene difluoride membrane (Merck-Millipore), stained with Direct Blue, excised and transferred to separate wells in a flat bottom polypropylene 96-well plate (Corning Life Sciences). Spots were blocked with 1% (w/v) polyvinylpyrrolidone in 50% (v/v) aqueous methanol and washed with water. *N*-glycans were enzymatically released using 3.5 U *Flavobacterium meningosepticum N*-glycosidase F recombinantly expressed in *E. coli*

(Roche) in 20 ml $H_2O$, 16 h at 37°C. The released *N*-glycans were hydroxylated using 100 mM aqueous $NH_4HCO_3$, pH 5, 1 h, 20°C, and reduced using 1 M $NaBH_4$ in 50 mM aqueous KOH, 3 h, 50°C. The reduction reaction was quenched using glacial acetic acid. Dual desalting of the reduced *N*-glycans was performed using first strong cation exchange resin (where the *N*-glycans are not retained) and then porous graphitised carbon (PGC) (where *N*-glycans are retained) packed as micro-columns on top of $C_{18}$ discs in P10 solid-phase extraction (SPE) formats. The *N*-glycans were eluted from the PGC SPE micro-columns using 0.05% trifluoroacetic acid (TFA) in 40% acetonitrile (ACN) and 59.95% $H_2O$ (v/v/v), dried and reconstituted in 20 µl $H_2O$, centrifuged at 14,000 x g, 10 min, 4°C and transferred into high recovery glass vials (Waters) for LC-MS/MS analysis. Bovine fetuin was included as a reference glycoprotein to ensure efficient *N*-glycan release and clean-up and to benchmark the LC-MS/MS instrument performance.

*N-glycan profile by PGC-LC-MS/MS*

Detached *N*-glycan mixtures from each granule sample were injected on a heated (50°C) PGC-LC capillary column (Hypercarb KAPPA, particle/pore size 5 µm/200 Å, column length 100 mm, inner diameter 180 µm, Thermo Fisher Scientific, Australia). Separation of *N*-glycans was carried out over a linear gradient of 0–45% (v/v) ACN (solvent B) in 10 mM aqueous $NH_4HCO_3$ (solvent A) for 75 min at a constant flow rate of 4 µl/min delivered by a Dionex Ultimate-3000 HPLC system (Sydney, Australia). *N*-glycans were analysed in negative polarity mode using electrospray ionisation (ESI) on a Velos Pro linear ion trap mass spectrometer (Thermo Fisher Scientific) with a MS1 full scan acquisition range of *m/z* 380-2,000. ESI was performed using an electrospray voltage of 3.2 kV, a nitrogen drying gas flow of 7 l/min at 275°C, and a nitrogen-based nebuliser pressure of 12 psi. An isolation window of 2 *m/z*, a minimum signal of 300 counts and a maximum accumulation time of 30 ms were applied. Data-dependent acquisition was used to select the top nine most abundant precursors in each MS1 full scan spectrum for fragmentation using resonance activation collision-induced dissociation tandem mass spectrometry (CID–MS/MS) at a normalised collision energy (NCE) of 35%. All MS and MS/MS data were acquired in profile mode and dynamic exclusion was disabled. The mass spectrometer was tuned and calibrated using a tune mix (Thermo Fisher Scientific) prior to use.

*N-glycan data analysis*

Raw data were browsed and interrogated using Xcalibur v2.2 (Thermo Fisher Scientific). GlycoMod (http://www.expasy.ch/tools/glycomod) and GlycoWorkBench v2.1[4] (https://code.google.com/archive/p/glycoworkbench/) assisted the otherwise manual *de novo* glycan annotation as described[5]. Briefly, *N*-glycans were identified based on monoisotopic precursor mass, MS/MS fragmentation pattern and relative and absolute PGC-LC retention time. EIC-based glycan quantification was performed using Skyline v.19.1[6,7]. GlycoWorkBench v2.1[4] (https://code.google.com/archive/p/glycoworkbench/) was used to aid the manual annotation of glycan fragment spectra and to generate glycan cartoons. Glycomics raw data are available from GlycoPost (accession number GPST000315).

**Glycoproteome profiling**

*Protein digestion*

The protein extracts from the luminal fraction of the neutrophil granule preparations were precipitated overnight using acetone at -20°C. The pelleted proteins were resuspended in 100 mM aqueous ammonium bicarbonate to a final protein concentration of 0.1 µg/ml. Proteins were then reduced using 10 mM aqueous DTT, 56°C, 45 min, and alkylated using 25 mM aqueous IAA, 25°C, 30 min in the dark (all final concentrations). The alkylation reactions were quenched using 30 mM aqueous DTT in 100 mM aqueous $NH_4HCO_3$, 25°C, 30 min. The proteins were then exhaustively digested using sequencing-grade modified porcine trypsin (Promega, Australia) at a 1:30 enzyme:substrate ratio (w/w) in 100 mM aqueous $NH_4HCO_3$, pH 8.4, 37°C, 16 h. Digestions were stopped by acidification with 1% (v/v) aqueous formic acid (FA, final concentration). The resulting peptide mixtures were desalted using C18-SPE micro-columns (Merck-Millipore, Australia). The desalted peptide mixtures were dried and redissolved in 10 µl 0.1% (v/v) aqueous FA for LC-MS/MS.

*Glycopeptide enrichment*

Glycopeptide enrichment was performed as described[8]. In short, dried peptides were dissolved in 10 µl loading solvent consisting of 80% ACN:1% TFA:19% $H_2O$ (v/v/v), and then applied to custom-made hydrophilic interaction liquid chromatography (HILIC) SPE micro-columns packed to a column height of 22 mm in GELoader tips (Eppendorf, Australia) with zwitterionic HILIC resin (ZIC-HILIC, 10 µm particle size, 200 Å pore size, kindly donated by Merck KGaA, Darmstadt, Germany) on top of C8 Empore SPE discs (Sigma-Aldrich). The micro-columns were equilibrated in 50 µl loading solvent. Approximately 70% of each peptide sample was loaded and reloaded in successive rounds before the micro-columns were washed twice with 50 µl loading solvent. The retained glycopeptides were eluted twice with 50 µl 1% (v/v) aqueous TFA and then with 50 µl loading solvent to ensure complete glycopeptide elution. The eluted fractions were combined and used for LC-MS/MS analysis (enriched glycopeptide fractions). The remaining 30% of each peptide sample was used directly for LC-MS/MS analysis (unenriched peptide fractions) after desalting on C18 SPE micro-columns packed in stage tips (Merck-Millipore).

*Peptide and glycopeptide profiling by reversed-phase LC-MS/MS*

The unenriched peptide fractions were analysed using a Q-Exactive HF-X Hybrid Quadrupole-Orbitrap mass spectrometer (Thermo Scientific, Australia) coupled to an Easy nLC-1200 HPLC system (Thermo Scientific, Australia). The mass spectrometer was operated in positive ion polarity mode. Peptides were separated using reversed-phase C18 chromatography using a nano-LC column packed in-house with C18AQ resin (1.9 µm particle size, 120 Å pore size, 0.075 mm inner diameter x 500 mm length, Dr Maisch HPLC GmbH, Germany). The mobile phases were 0.2% FA:19.8% $H_2O$:80% (v/v/v) ACN (solvent B) and 0.1% (v/v) aqueous FA (solvent A). The linear gradient of solvent B increased from 5-30% over 120 min, 30-60% over 5 min, 60-95% over 5 min and was kept for 5 min at 95% B. The flow rate was kept constant at 300 nl/min. The MS1 acquisition range was *m/z* 350-1,650 at a resolution of 60,000 (measured at *m/z* 200). The MS1 automatic gain control (AGC) target was 3 x $10^6$ with a maximum injection time of 50 ms. Data-dependent acquisition was used to select the fifteen most abundant precursors in each MS1 scan for MS/MS using higher-energy collisional dissociation (HCD) at 27% NCE. Precursor isolation windows of *m/z* 1.4 were used. The MS/MS AGC target was 1 x $10^5$, the maximum injection time was 25 ms and the resolution was 15,000 (measured at *m/z* 200). Already selected precursors were dynamically excluded for 20 s. Unassigned, singly charged precursors and highly charged precursors (Z > 8) were excluded for MS/MS.

The enriched glycopeptide fractions were analysed using an Orbitrap Fusion (Thermo Scientific) coupled to a Dionex 3500RS HPLC system. Glycopeptides were separated using reversed-phase C18 chromatography using a heated (45°C) nano-LC column packed in-house with C18AQ resin (1.9 µm particle size, 120 Å pore size, 0.075 mm inner diameter x 500 mm length, Dr Maisch HPLC GmbH, Germany) with a gradient of 5 –33% solvent B over 115 min at 300 nl/min. MS1 scans were acquired in the range *m/z* 350 –1,800 (120,000 resolution, 4 x $10^5$ AGC, 50 ms injection time) followed by HCD-MS/MS data-dependent acquisition of the seven most intense precursor ions featuring the highest charge state. Fragments were detected in the Orbitrap (30,000 resolution, 2 x $10^5$ AGC, 120 ms injection time, 40 NCE, 2 *m/z* quadrupole isolation width). Already selected precursors were dynamically excluded for 60 s. The acquisition strategy included a product ion triggered re-isolation of precursor ions if glycopeptide-specific HexNAc oxonium ions (*m/z* 138.0545, 204.0867, 366.1396) were detected amongst the top 20 fragment ions in each HCD-MS/MS spectrum. The re-isolated precursor ions were subjected to both ETD- and CID-MS/MS analysis. ETD-MS/MS fragments were detected in the Orbitrap (30,000 resolution, 2 x $10^5$ AGC, 120 ms injection time, 2 *m/z* quadrupole isolation width, calibrated charge-dependent ETD reaction times enabled) and CID-MS/MS fragments were detected in the ion trap (rapid scan rate, 2 x $10^4$ AGC, 70 ms injection time, 30% NCE, 2 *m/z* quadrupole isolation width).

Proteomics and glycoproteomics LC-MS/MS data were acquired from four biological replicates from each of the Az, Sp and Ge granule populations and three biological replicates for the Se fraction due to the loss of a biological replicate during sample build-up.

*Proteomics data analysis*

For protein identification and quantification, the raw files were imported into MaxQuant v1.6.10.43[9]. The Andromeda search engine was used to search the HCD-MS/MS data against the reviewed UniProtKB Human Protein Database (released December 2019; 20,364 entries) with a precursor and product ion mass tolerance of 4.5 ppm and 20 ppm, respectively. Carbamidomethylation of cysteine (57.021 Da) was set as a fixed modification. Oxidation of methionine (15.994 Da) and protein N-terminal acetylation (42.010 Da) were selected as variable modifications. All identifications were filtered to 1% protein FDR using a conventional decoy approach. For label-free AUC-based quantification, the ‘match between runs’ feature of MaxQuant was enabled with a 0.7 min match time window and 20 min alignment time window. Protein abundance was calculated based on the normalised protein intensity (LFQ intensity)[10].

*Glycopeptide identification and quantification*

Glycopeptides were identified and quantified from the HCD-MS/MS data using Byonic v4.5.2 (Protein Metrics Inc) operated as a node in Proteome Discoverer v.2.5. Search parameters included the use of i) a human proteome database (UniProtKB, 20,416 entries, downloaded August 2019), ii) a glycomics-informed *N*-glycan database comprising the 47 *N*-glycan compositions identified with glycomics of the same samples, iii) a search strategy permitting up to one *N*-glycan per peptide as a “rare” variable modification, semi-trypsin cleavage patterns with a maximum of two missed tryptic cleavages per peptide, up to 10/20 ppm mass tolerance of precursor/product ions from expected values, and up to one Met oxidation (+15.994 Da) per peptide (variable “common” modification), and by using monoisotopic correction (error check = +/- floor (mass in Da / 4,000)), and a decoy and a default contaminant database available in Byonic. Glycopeptide identifications were filtered to FDR < 1% (high confidence as determined using Proteome Discoverer 2.5). Glycopeptides identified with lower confidence, and those found in the decoy and contaminant database were excluded. Quantitation of (glyco)peptides was based on the area-under-the-curve of extracted ion chromatograms of precursor ions as determined by the Minora Feature Detector with the minimum trace length set to 5 and maximum delta retention time of isotope pattern multiplets of 0.2 min within Proteome Discoverer v2.5. The feature mapper tool was enabled to allow chromatographic alignment of signals and extraction of precursor area based on a matched retention time and *m/z* within a time and mass tolerance window determined automatically by the software. Minimum S/N thresholds were enabled. The matching CID- and ETD-MS/MS spectra were used to manually confirm the glycopeptide identity including glycan structure and glycosylation sites of glycopeptides of particular interest but were neither used for quantitation nor comparison across conditions. Relative glycopeptide quantification was performed by dividing the area of each peptide by the summed area of all peptides within each sample. The quantification of the site-specific glycoform distribution was performed by dividing the area of each glycoform by the summed area of all glycoforms identified within the same site from the same glycoprotein. For comparisons of the glycan composition/glycan class distribution across granules and fractions, glycoPSM counts were used and normalised against the total glycoPSM counts from each sample. The relative glycosylation efficiency (measure of relative glycan occupancy) was established by dividing the summed area of all glycopeptides by the summed area of all non-glycosylated peptides from the same samples. The glycoproteomics and proteomics raw files are available from the ProteomeXchange consortium via PRIDE accession numbers PXD039387 and PXD021131, respectively.

**Interrogation of available proteomics and transcriptomics resources**

Proteomics data of neutrophil precursors from four discrete maturation stages isolated by fluorescence-activated cell sorting of myeloid progenitor cells derived from the bone marrow of four healthy donors were obtained from the ProteomeXchange consortium via the PRIDE partner (data accession number PXD013785)[11]. The investigated maturation stages include the promyelocytic and myelocytic stage (collectively referred to as P/M), metamyelocytic stage (MM), immature neutrophils with band-formed nuclei (BN), and mature neutrophils with segmented nuclei (SN). Circulating (mature, resting) neutrophils (polymorphonuclear cells, PMN) derived from blood from the same four donors were also investigated. Multiple LC-MS/MS datasets (technical replicates) were available for each of these cell populations. For our study, a single technical replicate from each maturation stage from each of the four donors was interrogated given the existence of biological replicates. The LC-MS/MS data were searched using Byonic v4.5.2 (Protein Metrics Inc) operated as a node in Proteome Discoverer v.2.5. Search parameters were the same as used for the granule glycoproteomics search (see above). For glycoproteome quantification during neutrophil maturation, glycoPSM counts were used with a subtraction approach involving subtraction of glycoproteins identified in the maturation stage(s) prior to the specific maturation stage being assessed (hence only considering newly expressed glycoproteins at each stage).

RNAseq transcriptome data of neutrophils derived from blood and maturing neutrophils (P/M, MM, BN, SN) isolated from bone marrow of three healthy donors was obtained from Grassi et al.[12] and available at https://blueprint.haem.cam.ac.uk/neutrodiff. Proteomics and related RNA-seq data of the OST enzyme complex were obtained from the Supplementary Table S1 from Hoogendijk et al.[11]. For the transcriptome comparison between P/M vs MM, log2 fpkm values were obtained from a list of 3,456 genes available in the Supplementary Table S1 from Hoogendijk et al.[11]. An additional list of log2 fpkm values from 28 genes related to the *N*-glycosylation machinery were manually obtained from the source data available at https://blueprint.haem.cam.ac.uk/neutrodiff and added to the transcriptome data. Annotation of the genes as glycoproteins was performed using the glycoproteomics data acquired in this study. Significance was tested using t-test and multiple test correction and the data visualised as volcano plots. For the OST transcriptome and proteome comparison, transcript values (log2 fpkm) and protein level values (log 2 LFQ) across all four maturation stages (P/M, MM, BN and SN) were obtained from Supplementary Table S1 from Hoogendijk et al.[11]. Data were represented using scatter plots and tested for significance between P/M vs all the other stages using t-tests.

**Quantitation and statistical analyses**

Statistical significance was assessed using unpaired two-tailed Student's t-tests or ANOVA followed by Tukey test for multiple comparison with FDR < 0.05 as the confidence threshold. For transcriptomics and glycomics data analyses, the *p* value was corrected for multiple test comparison and values below 0.05 were considered significant. GraphPad Prism v9.4.1 (Dotmatics) and Perseus v2.0.7.0[13] were used for the statistical analysis. Correlation analysis was carried out using Pearson correlation coefficient using Excel. PCA analysis was performed using Metaboanalyst v. 5.0[14].

**Data sharing statement**

To provide an accessible resource featuring comprehensive spatiotemporal information of the neutrophil *N*-glycoproteome for community interrogation and hypothesis generation/testing, all generated LC-MS/MS datasets, search output and metadata have been deposited in public repositories. Specifically, glyco/proteomics LC-MS/MS raw data, search files (.msf) and search results (.pdResult) have been deposited to the ProteomeXchange Consortium via the PRIDE repository and are publicly available as of the date of publication (Accession numbers PRIDE ID: PXD039387 and PXD021131). Glycomics LC-MS/MS raw data have been deposited to the GlycoPOST repository (Accession numbers GlycoPOST ID: GPST000315).
This paper analyses existing publicly available data including proteomics (PRIDE ID: PXD013785)[11] and transcriptomics (https://blueprint.haem.cam.ac.uk/neutrodiff/)[12] data collected from developing myeloid progenitors.

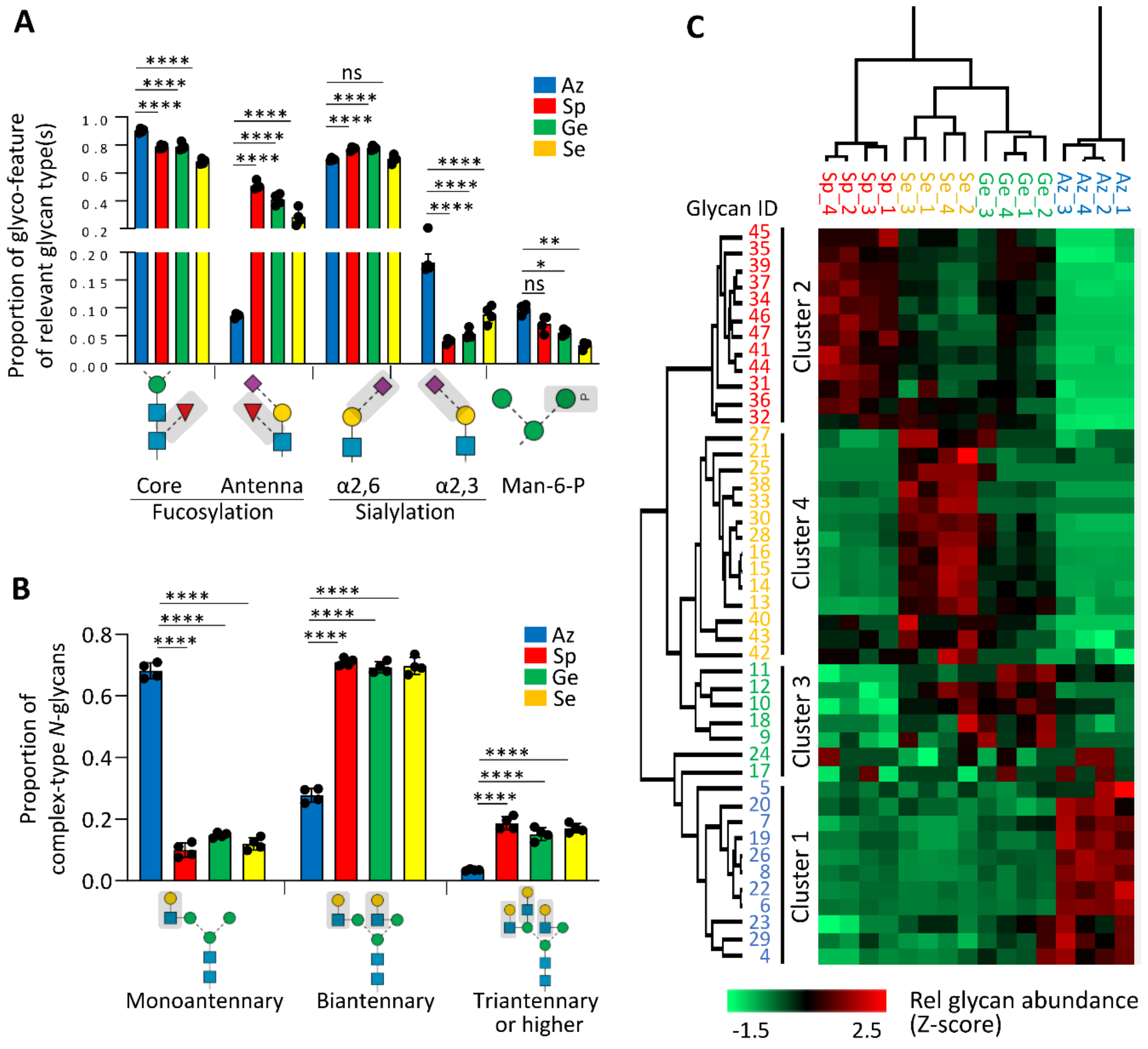


**Supplementary Figure S1. Granule-specific *N*-glycome features in resting neutrophils.** Distribution patterns of key *N*-glycan features across granules including **A**) core/antenna fucosylation, α2,3-/α2,6-sialylation and mannose-6-phosphate (M6P) modifications and **B**) antennary branching. For panel A and B, data points are plotted as mean and SD relative abundance of each glycan feature against the relevant *N*-glycan type(s) that has the potential to be modified by that feature (n = 4 biological replicates, ANOVA test, ***p* < 0.01, *****p* < 0.0001, ns, not significant). **C**) Unsupervised clustering and heatmap of *N*-glycomics data illustrating four major clusters representing the four investigated neutrophil granule populations (full view, expanded from condensed view in **Figure 2F**). See **Supplementary Table S2** for overview of *N*-glycan identifiers and tabulated quantitative *N*-glycomics data. See **Figure 1** for key.

**A**

| Reiding et al | GlycoPSMs | Glycopeptides | Glycoproteins |
|---|---|---|---|
| Enriched | 9,324 | 1,963 | 140 |
| Unenriched | 3,753 | 470 | 42 |
| Total | 13,077 | 2,318 | 152 |

| Kawahara et al Granule data | GlycoPSMs | Glycopeptides | Glycoproteins |
|---|---|---|---|
| Enriched | 23,191 | 4,554 | 332 |
| Unenriched | 4,472 | 716 | 115 |
| Total | 27,663 | 4,772 | 352 |

| Maturation data | GlycoPSMs | Glycopeptides | Glycoproteins |
|---|---|---|---|
| Unenriched | 11,928 | 2,276 | 1,079 |

**B**

**C**

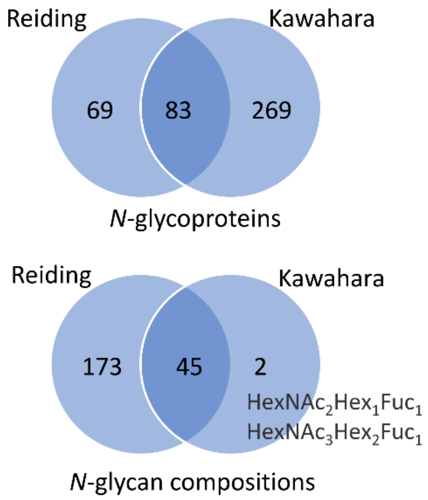


**Supplementary Figure S2.** Neutrophil *N*-glycoproteome coverage reported in this study relative to reference study. **A**) Overview of the *N*-glycoproteome coverage including the total *N*-glycoPSM count, unique *N*-glycopeptides (unique protein, site, glycan) and source *N*-glycoproteins identified (top table) across unfractionated neutrophil samples in a reference study (Reiding et al.)[15], (middle table) across granules with and without glycopeptide enrichment in this study (Kawahara et al.) and (bottom table) during neutrophil granulopoiesis. For the latter, maturation data were obtained by reinterrogation of proteomics resources available via PRIDE accession PXD013785 from Hoogendijk et al.[11]. Comparison of **B**) source *N*-glycoproteins and **C**) *N*-glycan compositions reported by a reference study (Reiding)[15] and this study (Kawahara) based on neutrophil glycoproteomics data.

A

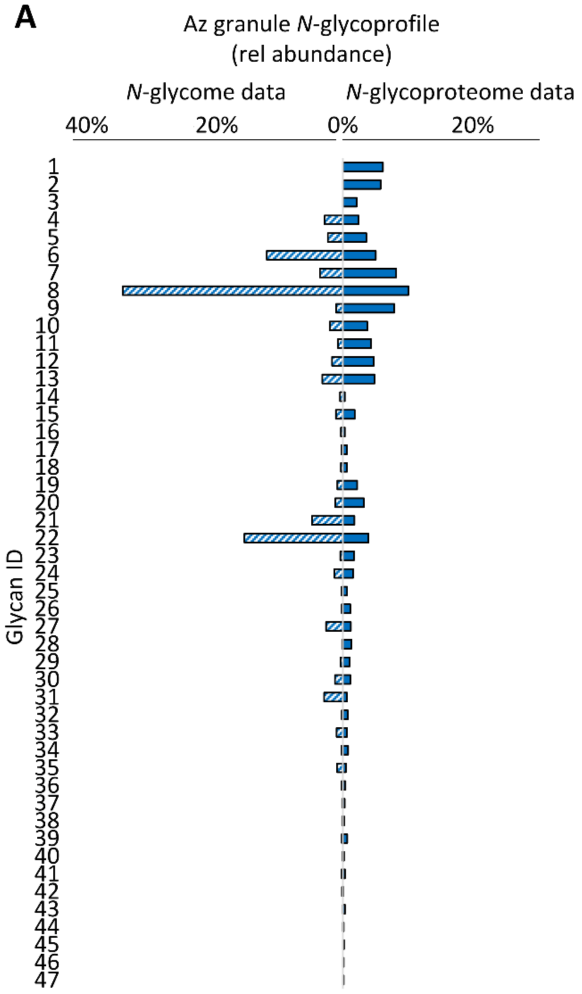


C

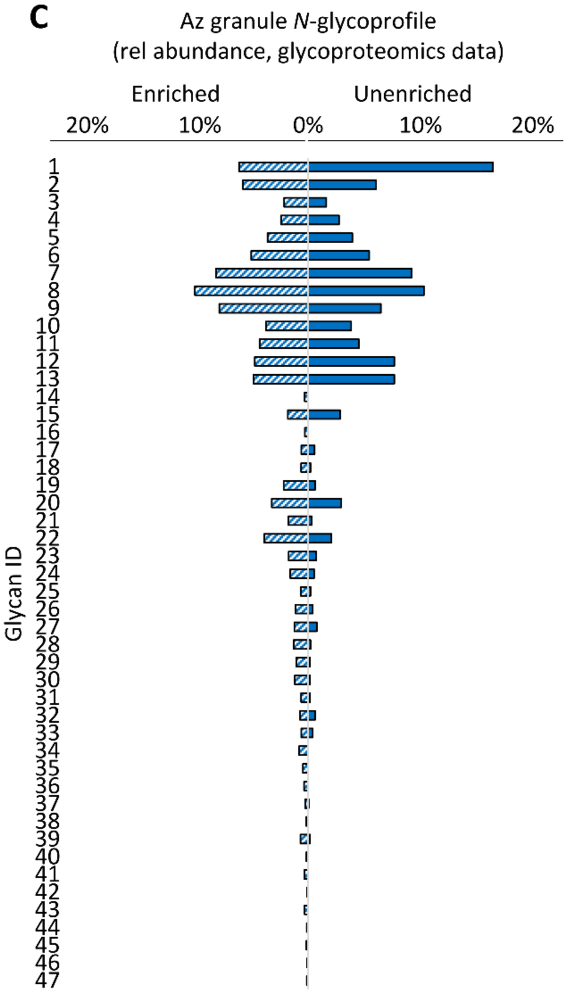


**B**

| Pearson correlation (R) | Az | Sp | Ge | Se |
|---|---|---|---|---|
| Glycomics vs glycoproteomics | 0.588 | 0.505 | 0.529 | 0.419 |

**D**

| Pearson correlation (R) | Az | Sp | Ge | Se |
|---|---|---|---|---|
| Enriched vs unenriched | 0.875 | 0.649 | 0.789 | 0.906 |

**Supplementary Figure S3. Correlation between glycomics and glycoproteomics data**. **A**) Quantitative comparison of *N*-glycan distribution patterns identified from glycomics and glycoproteomics data of the Az granules. See **Supplementary Table S2** for overview of *N*-glycan identifiers and *N*-glycomics data. See **Supplementary Table S3** for *N*-glycoproteomics data. **B**) Pearson correlation (R) comparing the distribution of *N*-glycans identified with glycomics and glycoproteomics across the four investigated neutrophil granules. **C**) Quantitative comparison of *N*-glycan distribution patterns identified from glycoproteomics data acquired with and without glycopeptide enrichment of the Az granules (**Supplementary Table S3**). **D**) Pearson correlation (R) comparing the distribution of *N*-glycans identified in glycoproteomics data acquired with and without glycopeptide enrichment across the neutrophil granules.

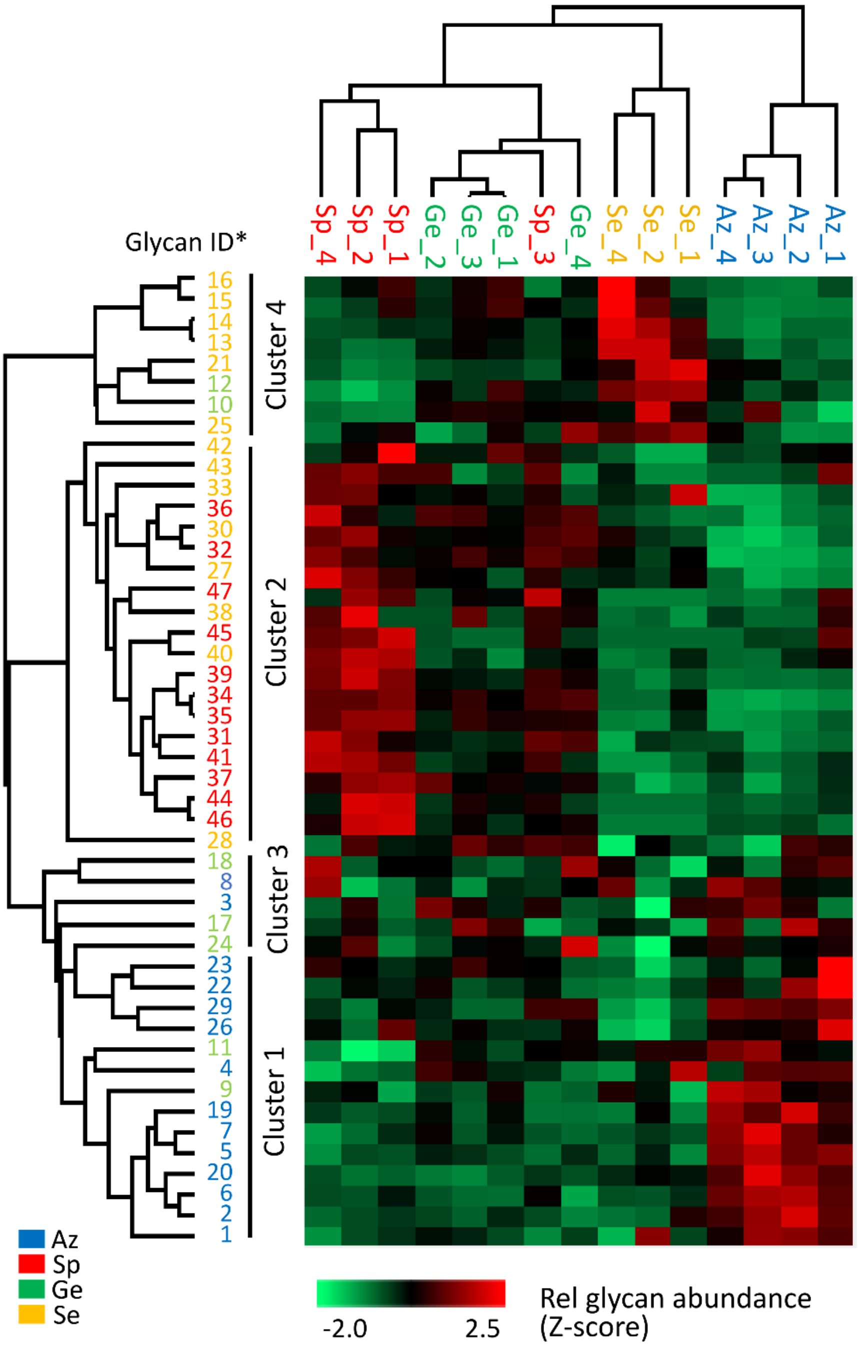


**Supplementary Figure S4. Similar glycoproteomics- and glycomics-guided clustering patterns confirms granule-specific *N*-glycosylation in resting neutrophils.** Unsupervised clustering using Euclidean distance and average linkage of Z-score transformed relative abundance data of *N*-glycan compositions identified in the glycoproteomics data generated from the neutrophil granules. See **Supplementary Table S2** for overview of *N*-glycan identifiers. Row clusters are colour coded according to the *N*-glycomics-guided clustering (see **Figure 2F**) to illustrate the similarity between the two cluster analyses informed by the two different datasets.
